# What is memorable across the Pacific Ocean shows evidence of prior learning on the memorability of new visual inputs

**DOI:** 10.1101/2025.07.23.666320

**Authors:** Haoyu Chen, Benchi Wang, Geoffrey F Woodman

## Abstract

It has been proposed that certain objects in our environment are inherently memorable due to fundamental similarities in how the human brain stores this visual information. Here we tested this hypothesis by having two groups of participants from opposite sides of the planet remember pictures of objects shown individually from several seconds each. We found that the specific objects that were more memorable varied across groups, with fine-grained analysis of the nonoverlapping objects indicating that participants have greater facility storing objects that are familiar to them as they occur in their cultural context. Thus, the present findings demonstrate that the memorability of a stimulus is jointly determined by its features and the learning history of the human brain in which it is being stored.

## Introduction

Memorability is the phenomenon in which certain stimuli are better remembered across observers, presumably due to stimulus-intrinsic properties that determine the likelihood of stimulus being successfully remembered (Bainbridge et al., 2013). It has captured the attention of memory and vision researchers due to its stability (Bainbridge, 2017), independence (Bainbridge, 2020), and generalizability (Bainbridge et al., 2013; Kramer et al., 2023; Madan, 2021). One of the dominant theoretical explanations of this heavily studied effect is that it reveals fundamental aspects of memory storage related to bottom-up visual feature processing that operates in every human brain regardless of individuals’ prior experiences (Bainbridge et al., 2013; Isola et al., 2014).

If we wanted the strongest experimental test the hypothesis that the stimuli determine what is memorable and not what the brain has stored previously, then we would raise separate groups of animals in different environments, with different objects and different statistics, perhaps in cages on opposite sides of the Earth. Then we could see if learning history changed which objects were memorable. Here we did the next best thing, which was to study memorability in separate groups of humans raised on opposite sides of the Earth (i.e., China versus the United States of America, we will use just America henceforth). If our experiences shape how well our brains can accommodate new memory representations, as theories of long-term memory would suggest (Shiffrin and Steyvers 1997, Polyn, Norman et al. 2009), then we should see what is memorable changes between the groups of participants raised in different environments. Although internet homogenization precludes perfectly isolated groups, the China vs. America comparison achieves a geographic and cultural maximum: divergent pre-digital visual environments and cultural practices generate residual differences in early-life experience. Using human subjects, this is one of the strongest possible tests of experience-dependent memorability.

Studies of memory performance across large groups of participants have shown that images possess intrinsic memorability, defined as the likelihood of stimulus being remembered for any given person (Bainbridge et al., 2013; Isola et al., 2014; Khosla et al., 2015; Mancas & Le Meur, 2013). Crucially, this operational definition conceptualizes memorability as an intrinsic property of the stimulus itself, thereby postulating a testable hypothesis: If memorability is fundamentally inherent to the stimulus and remains consistent across individuals regardless of their prior experiences, it should logically exhibit high consistency across cultural contexts.

Our alternative hypothesis proposes experience-dependent memorability. Given that individuals from distinct cultural backgrounds possess divergent experiences, with culture acting as a filter, they simply experience different things, both as physical objects or experienced through media technologies (Connerton, 1989; Heersmink, 2023; Hobsbawm & Ranger, 2012). Based on this hypothesis, things that are memorable should be culture specific. However, previous research also highlights the generalization and consistency of memorability. Memorability exists in a wide range of stimuli including sounds, images, and words (Isola et al., 2014; Revsine et al., 2025; Xie et al., 2020), and remain consistent even across human and monkeys: memorability estimated by models designed to predict image memorability in humans correlate with monkeys’ memory performance for images (Jaegle et al., 2019). This cross-species consistency in memorability suggests that it may constitute a general cognitive mechanism rooted in shared neurobiological substrates, thereby implying potential invariance across cultures. However, so far, no studies have systematically investigated the cross-cultural consistency of memorability. Existing memorability research predominantly samples Western populations, creating a methodological blind spot regarding cultural variability. This monocultural focus limits our understanding of how memorability interacts with the cultural context in which the observer is embedded.

We took advantage of a neural network model developed by Kramer et al. (2023). This deep convolutional neural network (DCNN) model was used to identify a 49-dimensional feature space that explained 38.52% of variance in object memorability in a stimulus set. These dimensions—derived through data-driven computational modeling of 1,854 real-world object images in a previous study—were optimized to maximize explanatory power while maintaining psychological validity and categorical comprehensiveness (Hebart et al., 2020). Here we will use this model to identify cultural variations in the factors influencing object memorability across different cultural contexts, when such differences arise.

In summary, we tested subjects’ memory while looking for influences of culture on memorability. First, we used the same methods and stimuli as Saito et al. (2023) to obtain memorability scores from a sample of Chinese participants (see Methods for details). Next, we conducted correlation analyses to determine the extent of cross-cultural consistency in memorability. Our results revealed a strong correlation, suggesting significant cross-cultural consistency. Next, we implement the DCNN-based modeling to identify culture-specific determinants of memorability, with the model identifying specific semantic categories of objects that our Chinese versus American participants found easier to remember.

## Methods

### Participants

One hundred and twenty participants (*M*_age_ = 23.9, *SD* = 3.9, *n*_female_=55) were recruited from the online platform NAODAO (https://www.naodao.com) in China. This online study was approved by the local ethics committee (2020-3-013), and all participants provided informed consent before testing. Additionally, we incorporated data from Experiment 1A in Saito et al. (2023), which included 120 participants resided in the United States and Canada (*M*_age_= 24.2, SD = 3.9, *n*_female_ = 61).

### Stimuli

We used the same stimuli as Saito et al. (2023), consisting of 600 images of randomly chosen real-world objects from an existing database (Brady et al., 2008). The stimulus set was randomly divided into four subsets, each containing 150 images. For each participant, one subset was designated as “old” pictures, while another was used as “new” pictures. Each subset was assigned as “old” and “new” for 30 participants, respectively.

### Procedure

Participants began with an encoding task (Fig. 1a) consisting of 150 trials. Each trial started with a 500 ms fixation cross, followed by an image presented centrally for 1000 ms. After a 500 ms blank interval, a question appeared alongside a 6-point Likert scale, asking participants, “Are you going to remember the picture you just saw?” Participants responded using the scale (1 = definitely yes, 2 = probably yes, 3 = maybe yes, 4 = maybe no, 5 = probably no, 6 = definitely no). The Likert scale remained on the screen until a response was provided. Each participant viewed 150 new images, with the presentation order randomized across individuals.

**Figure 1.**
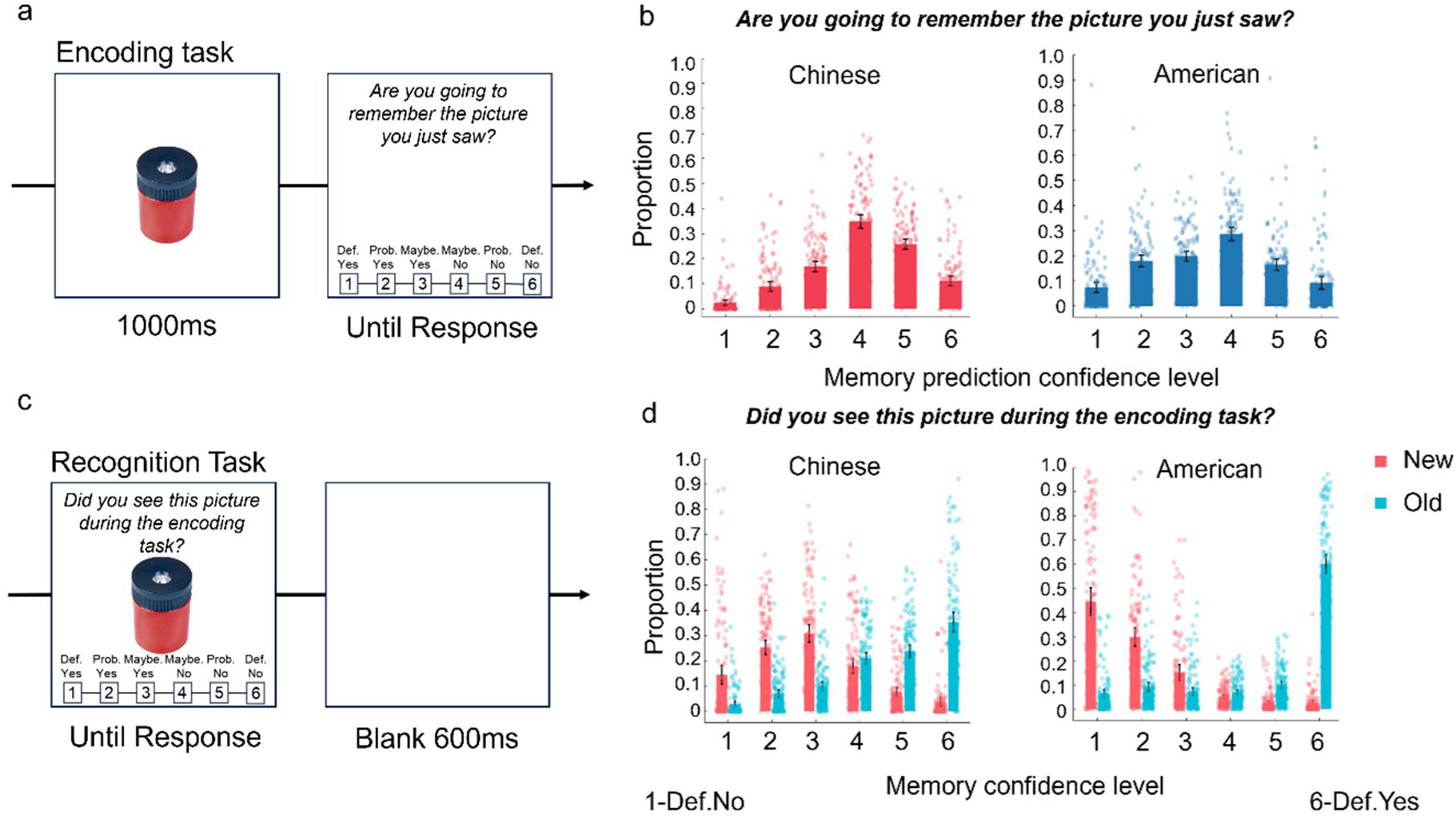
Experiment procedures, the distributions of memorability predictions, and response confidence across cultures. a) The Encoding Task shown with a randomly selected stimulus. Participants completed 150 trials. In each trial, after presenting a new image for 1000 ms, a question (“Are you going to remember the picture you just saw?“) and a 6-point Likert scale appeared on the screen. Participants responded by selecting one of six options (1 = definitely no, 6 = definitely yes). b) Results of participants’ predictions of memorability. The mean proportions of participants’ prediction confidence levels for Chinese participants (2.43%, 8.88%, 16.89%, 34.86%, 25.79%, and 11.15%) and American participants (7.51%, 18.01%, 19.92%, 28.69%, 16.52%, and 9.35%), from confidence level 1 to 6, respectively. c) The Recognition Task shown with the same example stimulus as in a). It consisted of 300 trials. Each trial began with a test stimulus presented at the center of the screen, accompanied by a question (“Did you see this picture during the encoding task?“) displayed above the image. Below the image, a 6-point Likert scale (1 = definitely no, 6 = definitely yes) allowed participants to make their recognition judgments. d) Memorability scores. For Chinese participants, the mean proportions of confidence levels (1–6) were 2.88%, 7.07%, 9.96%, 21.03%, 23.74%, and 35.32%, respectively for old images, and 14.45%, 25.26%, 30.84%, 17.99%, 7.72%, and 3.74%, respectively for new images. For American participants, the mean proportions of confidence levels (1– 6) were 6.46%, 9.21%, 7.19%, 7.02%, 10.02%, and 60.11%, respectively for old images, and 44.50%, 29.84%, 15.14%, 5.13%, 2.71%, and 2.68%, respectively for new images. A score of 1 represents “definitely no” and 6 represents “definitely yes“.

After completing the encoding task, participants proceeded to a recognition task (Fig. 1c) comprising 300 trials. Each trial presented a test image centrally, accompanied by the question, “Did you see this picture during the encoding task?” displayed above the stimulus, with the same 6-point Likert scale below. The Likert scale remained on the screen until a response was provided. Following each recognition judgment, the screen remained blank for a 600 ms inter-trial interval. The recognition task included 150 old images (seen during encoding) and 150 new images, presented in random order.

### Data Analysis

#### Memorability Scores

Memorability scores for each stimulus were calculated by subtracting the average recognition response when the stimulus was new (average “new” response) from the average recognition response when it was old (average “old” response). To enhance the interpretability of these scores, participants’ responses were reverse-coded so that higher values corresponded to greater confidence in remembering (e.g., 6 = definitely yes) and lower values to less confidence (e.g., 1 = definitely no). Consequently, more positive memorability scores indicated higher memorability, while less positive scores indicated lower memorability.

Participants’ ability to predict memorability was assessed by averaging the predicted memory responses for each stimulus (with each stimulus receiving 30 estimations). Similarly, these predictions were reverse-coded to ensure that higher values reflected higher expected memorability (e.g., 6 = definitely yes), and lower values indicated lower expected memorability (e.g., 1 = definitely no).

#### Object-dimension scores

To determine object-dimension scores for our stimuli, we employed a DCNN-based (Deep Convolutional Neural Networks) prediction method similar to that used in Kramer et al. (2023). Specifically, we utilized the THINGS dataset, which provides 49 object-dimension scores for 1,854 real-world images (Hebart, 2020), to train Elastic Net regression models. These models were then used to generate 49-dimension predictions for our stimulus set of 600 images (Brady objects). That is, we trained a separate Elastic Net regression model for each dimension, using activations from the penultimate layer of the CLIP Vision Transformer. This model has been demonstrated to exhibit highly human-like behavior across various tasks (Geirhos, 2021). Hyperparameters for the models were optimized using 10-fold cross-validation to maximize the fit for each dimension. The trained models achieved high predictive accuracy in most dimensions, with Pearson correlations between predicted and true dimension scores exceeding *r* > 0.9 for 13 dimensions, *r* > 0.8 for 33 dimensions, and *r* > 0.7 for 46 out of the 49 dimensions. Then, the regression weights from these models were applied to the DCNN representations of our stimulus set, generating 49-dimension scores for all 600 images. Each object dimension had its own label (e.g., *made of metal/artificial/hard* or *construction-related/physical work-related*). To enhance interpretability, we used Deepseek R1 to summarize these dimension labels into concise word pairs based on their names.

#### SVM-Based selection of important dimension

To identify key object dimensions distinguishing memorability between Chinese and American participants, we employed Support Vector Machine Recursive Feature Elimination (SVM-RFE) to select object dimensions that contribute to the classification between American memorable objects and Chinese memorable objects. We created SVMs (Support Vector Machines) that are supervised learning models designed to classify data by identifying the hyperplane that maximizes the margin between classes. By using kernel functions, SVMs can transform nonlinear problems into linearly separable ones in higher-dimensional space. The mathematical formulation of an SVM is as follow:

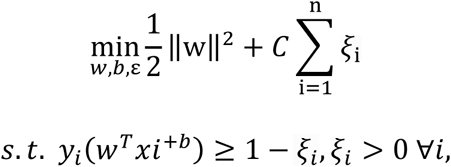

where *w* is the weight vector, *b* is the bias term, *ξ_i_* are slack variables allowing for margin violations in non-separable data, *C* is a regularization parameter balancing margin width and classification errors. The class label *y*_i_ of the *i*-th training sample is assigned as:

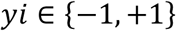

where y*_i_* = +1 represents the positive class, and y*i* = −1 represents the negative class.

The feature vector x*_i_* for the *i*-th training sample is represented as:

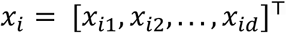

where *d* is the number of dimensions.

We employed the MATLAB function *fitcsvm.m*, and used 49 object dimensions of 143 objects (71 American-memorable and 72 Chinese-memorable objects; details can be found in the Results section) to train the SVM. To ensure robustness, the training process was repeated six times with 10-fold cross-validation. In each iteration, 129 images were used for training and 14 images were reserved for testing, resulting in a classification accuracy rate. To identify the most important dimensions, we calculated the average weights of each dimension in the SVM model and excluded the dimension with the least weight (indicating the least contribution to classification). The model was then retrained with the remaining dimensions. This process of exclusion and retraining was repeated iteratively until all dimensions were removed.

The key assumption underlying this procedure was that dimensions excluded later in the sequence were more important for classification. In total, the iterative process was repeated 49 times, each time removing one dimension, generating a sequence of 49 classification accuracy rates. These rates corresponded to the dimensions that remained at each step. To refine the analysis, we applied a 7-point moving average to smooth the classification accuracy rates and selected the dimensions corresponding to the peak accuracy rate as the most important dimensions. This approach allowed us to identify the subset of object dimensions that played the most significant role in distinguishing culturally specific memorability patterns.

#### Regression model analyses

To investigate how life and non-life dimensions influence memorability and its cultural specificity, we categorized object dimensions into three groups based on their labels. Life dimensions were defined as those related to life or body parts (e.g., *Animal/organic*, *Arms/skin*; see Fig. 3a). Non-life dimensions were defined as those unrelated to life or body parts (e.g., *Metal/artificial*, *Fire/heat*). Dimensions exhibiting both life and non-life traits or ambiguous characteristics (e.g., *Sky/flying*, which could refer to birds or aircraft) were categorized as the mixed one.

We conducted multiple regression analyses across both cultures to predict memorability. The top six life and non-life object dimensions most correlated with memorability were used as predictors separately, to match the number of object dimensions across conditions. We derived culture-specific memorability by fitting a linear regression model predicting one culture’s scores from another’s (the fitted line is shown in Fig. 2a, left). We subtracted the predicted values from the actual values, obtaining the residuals (i.e., the unpredicted part). These residuals represent culture-specific memorability. The underlying logic is that the shared component of memorability can be predicted by another culture’s scores, while the culture-specific component cannot. Then the culture-specific memorability was used as dependent variable to train the Life/Non-life multiple regression model.

**Figure 2.**
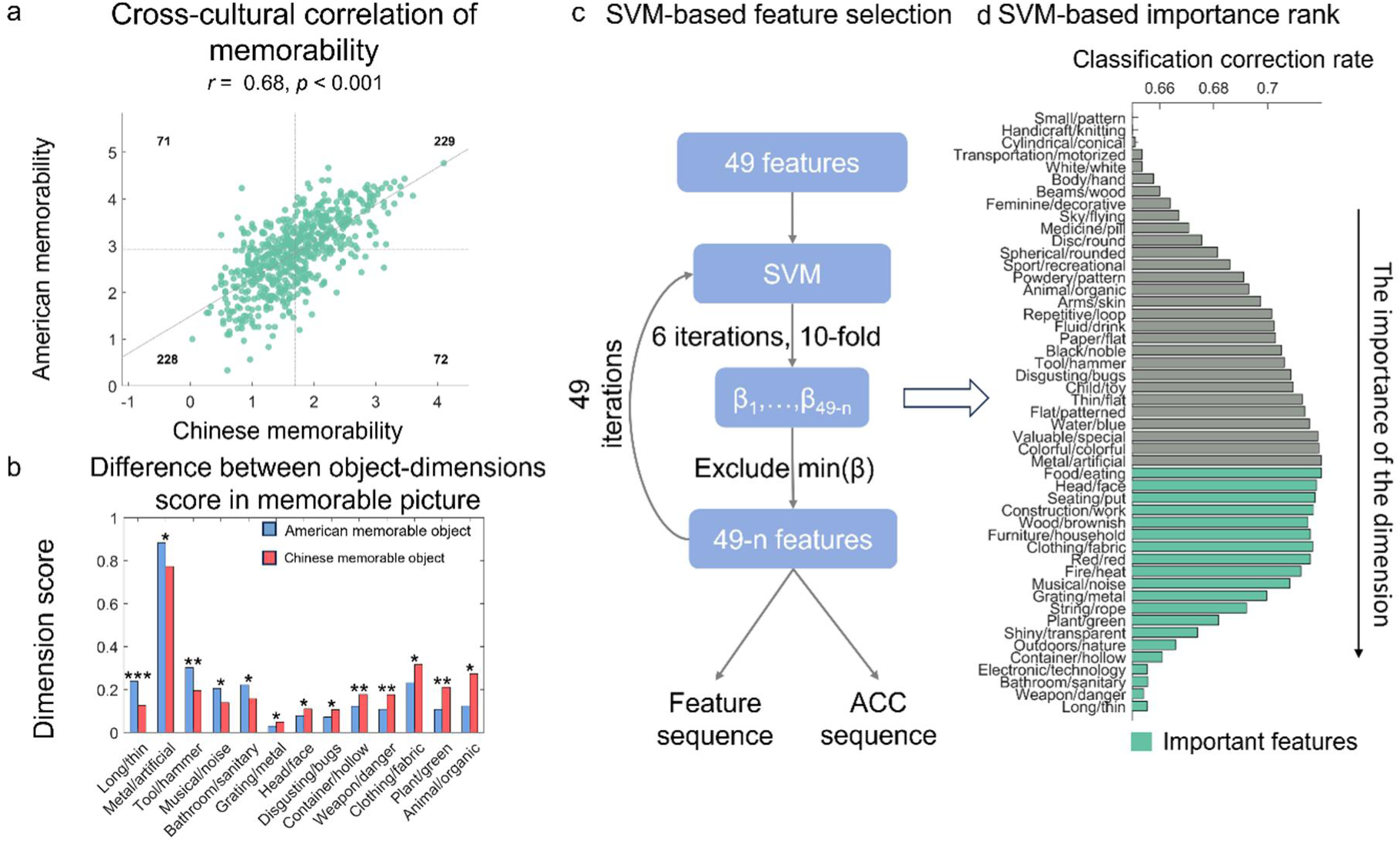
**a)** The cross-cultural correlation of memorability and its prediction, the scatter plot was split into four quadrants by the medium, the number in each quadrant correspond to the number of images belongs to this quadrant. b) Object dimensions that score significant differently between American memorable picture and Chinese memorable picture, significant dimensions, \**p* < .05, \*\**p* < .01, *** *p* < .001. c) The procedure of SVM-based feature selection. d) The rank of importance selected by SVM.

**Figure 3.**
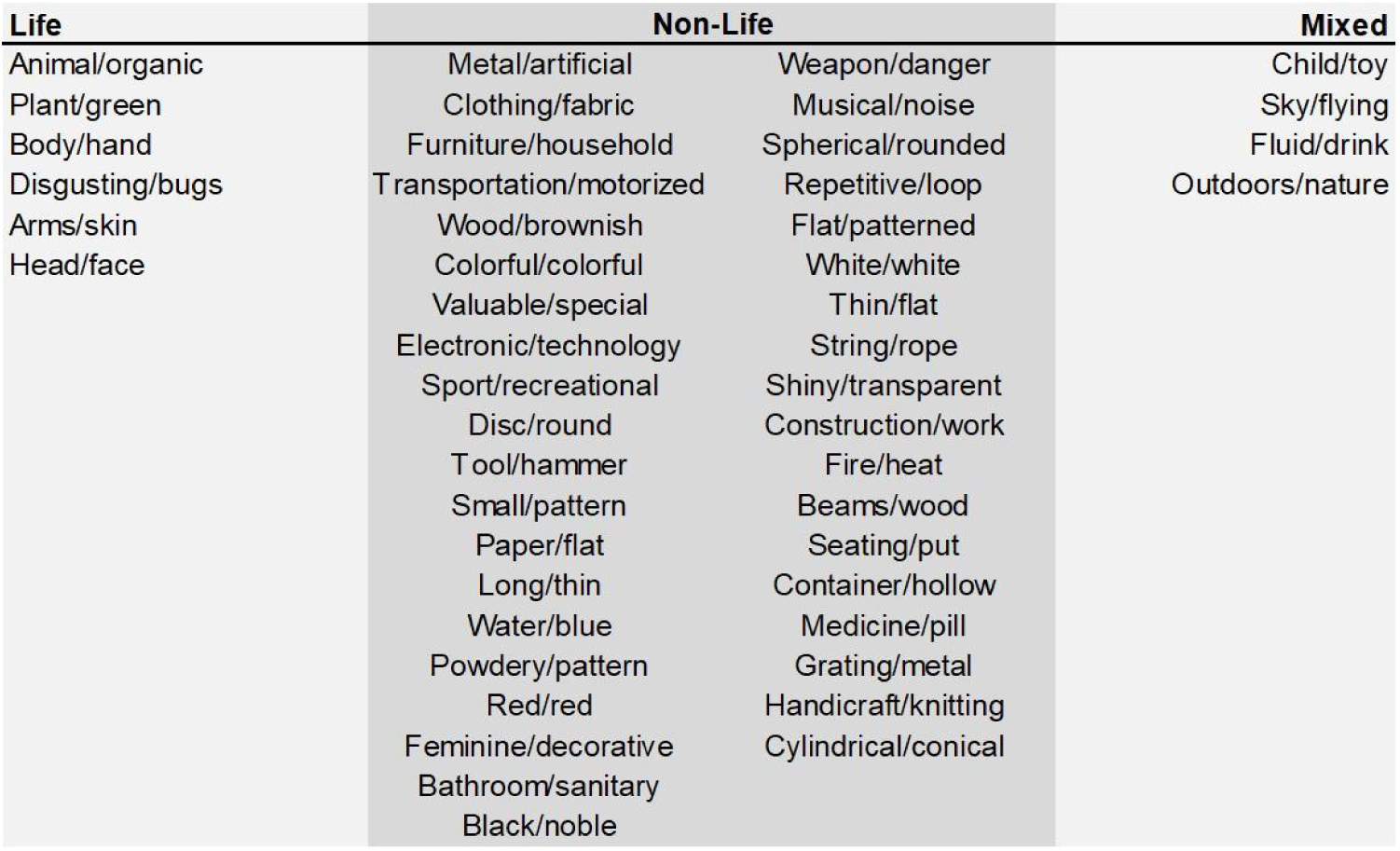
Categorization of THINGS object space dimensions across Life, Non-life, and mixed dimensions.

## Results

### Memorability predictions

For the prediction of memorability, across both cultures, the middle confidence level (i.e., the fourth one, “Maybe yes“) was most frequently selected, with both groups exhibiting the same general pattern (see Figure 1b). For memory judgment confidence levels, when presented with old objects (previously seen during encoding), participants from both cultures most often selected the highest confidence level (i.e., the sixth one, “Definitely yes“). The selection frequency for old objects declined progressively with each decrease in confidence level (Figure 1d), demonstrating that participants from both China and America exhibited similar patterns. We thus decided not to involve the prediction of memorability in further analysis.

Cultural differences emerged in confidence patterns for new objects (unseen during encoding). Chinese participants most frequently selected the middle confidence level (i.e., the third one, “Maybe no“), whereas American participants most often chose the first confidence level (“Definitely no“). Among American participants, the selection frequency for new objects decreased progressively as confidence levels increased (Figure 1d). These findings suggest that Chinese participants showed a higher tendency to judge new objects as old, while American participants demonstrated greater confidence in correctly identifying new objects. Despite these cultural differences, the overall patterns of memory judgments and their predictions were largely comparable across cultures.

### Cross-cultural consistency and inconsistency of memorability

A significant positive correlation was observed (*r* = 0.69, *p* < .001; see Figure 2a, left panel), indicating substantial consistency in memorability across cultures, using a Pearson correlation was calculated between memorability scores from Chinese and American participants. This shows that there is significant agreement across groups in terms of the memorability of the stimulus set, although clearly the correlation is not perfect, indicating group differences, as we discuss next.

Despite this observed cross-cultural consistency, an important question is whether cultural specificity exists in memorability. According to the experience-dependent memorability hypothesis (Connerton, 1989; Heersmink, 2023; Hobsbawm & Ranger, 2012) memory should be shaped by cultural context. To determine whether our data support such a conclusion, we sorted objects into four quadrants based on their median memorability ratings within each culture (Figure 2a). Quadrant 1 showed objects memorable in both cultures; quadrant 2 showed objects memorable for Americans but not for Chinese participants; quadrant 3 showed objects unmemorable in both cultures; quadrant 4 showed objects memorable for Chinese but not for American participants. This classification allows for an examination of cultural differences in memorability, complementing the observed cross-cultural consistency.

Using a deep convolutional neural network (DCNN)-based approach (Kramer et al., 2023, see Methods for details), 49-dimensional feature scores were extracted for all objects. A comparison of feature scores between American-memorable objects (Quadrant 2) and Chinese-memorable objects (Quadrant 4) revealed distinct patterns. American-memorable objects exhibited significantly higher scores than Chinese-memorable objects on five dimensions: *Long/thin*, *Metal/artificial*, *Tool/hammer*, *Musical/noise*, and *Bathroom/Sanitary* (all *p*s < .049, one-tailed).

Conversely, Chinese-memorable objects scored significantly higher than American-memorable objects on eight dimensions: *Grating/metal*, *Head/face*, *Disgusting/bugs*, *Container/hollow*, *Weapon/danger*, *Clothing/fabric*, *Plant/green*, and *Animal/organic* (all *p*s < .049, one-tailed; Figure 2b). These results suggested that feature dimensions scoring higher for American-memorable objects (e.g., *Tool/hammer*, *Metal/artificial*) reflect objective physical attributes, whereas dimensions scoring higher for Chinese-memorable objects (e.g., *Head/face*, *Plant/green*, *Animal/organic*) indicate stronger associations with culturally influenced life experience domains.

Although distinct cultural influences on memorability were observed when analyzing individual features, we further evaluated the stability of these results through a more objective analysis using Support Vector Machine Recursive Feature Elimination (SVM-RFE). This method, commonly applied in biological contexts for feature selection (Guan et al., 2024), was adapted here to identify the combination of object features most predictive of American-versus Chinese-memorable classifications (see Methods for details). SVM-RFE began with the full set of 49-dimensional feature vectors. Features were iteratively removed from the model based on their weights in the SVM, with the least important feature eliminated at each step. The sequence of elimination served as an inverse measure of feature importance—features retained until the final rounds were considered most critical for classification. Classification accuracy, measured at each iteration, determined the optimal subset of features. Results showed that accuracy peaked prior to the elimination of the Food/eating feature dimension (Figure 2d), corresponding to a subset of 20 dimensions. It includes: *Food/eating*, *Head/face*, *Seating/put*, *Construction/work*, *Wood/brownish*, *Furniture/household*, *Clothing/fabric*, *Red*, *Fire/heat*, *Musical/noise*, *Grating/metal*, *String/rope*, *Plant/green*, *Shiny/transparent*, *Outdoors/nature*, *Container/hollow*, *Electronic/technology*, *Bathroom/sanitary*, *Weapon/danger*, *Long/thin*.

These findings align closely with the results of direct feature comparisons across cultures, identifying nine overlapping features: *Long/thin*, *Musical/noise*, *Bathroom/sanitary*, *Grating/metal*, *Head/face*, *Container/hollow*, *Weapon/danger*, *Clothing/fabric*, *and Plant/green.* These features encompass both life-related attributes (e.g., *Head/face*, *Plant/green*) and objective physical characteristics (e.g., *Long/thin*, *Grating/metal*), highlighting their contributions to the cultural differences in memorability. Based on these observations, we propose that life-relevance dimensions (e.g., life-related versus non-life-related features) differentially shape memorability across cultures, potentially driving the observed divergence in memory patterns between American and Chinese participants.

### Model prediction of memorability based on life-relevance dimensions

We sorted object dimensions into Life/Non-life dimensions based on their labels. Life dimensions were defined as those related to life or body parts, such as *Animal/organic* or *Arms/skin* (Fig. 3a). Non-life dimensions were defined as those unrelated to life or body parts, such as *Metal/artificial* or *Fire/heat*. Dimensions exhibiting a mixture of life and non-life traits or ambiguous characteristics were categorized as the mixed one, such as *Sky/flying* (since flying could refer to a bird or an aircraft), which was not involved in future analysis.

Using these labeled dimensions, we analyzed the contributions of life and non-life dimensions to culture-specific memorability by training regression models. Culture-specific memorability was derived by regressing out the shared variance between the two cultures (see Methods for details), resulting in Chinese-specific and American-specific memorability scores. Separate regression analyses were conducted for life and non-life dimensions, with results summarized in Table 1. The non-life dimensions model significantly predicted both Chinese-specific memorability (variance explained: 12.05%) and American-specific memorability (variance explained: 6.04%). Similarly, the life dimensions model significantly predicted Chinese-specific memorability (variance explained: 12.61%). However, the life dimensions model did not significantly predict American-specific memorability (variance explained: 0.13%). These findings suggest that life dimensions have a stronger influence on Chinese-specific memorability, while they do not appear to play a significant role in shaping American-specific memorability.

**Table 1.** Summary of regression model results.

| Dependent Variable | Culture | Model | $R^2$ | $F$ | $p$ | Error |
| --- | --- | --- | --- | --- | --- | --- |
| Culture-specific memorability | Chinese | Life | 0.126 | 14.26 | $3.280 \times 10^{-15***}$ | 0.2018 |
| | | Non-Life | 0.121 | 13.54 | $1.995 \times 10^{-14***}$ | 0.2032 |
|  | American | Life | 0.001 | 0.13 | 0.993 | 0.3112 |
| | | Non-Life | 0.06 | 6.36 | $1.687 \times 10^{-6***}$ | 0.2928 |

\*\*\* $p < .001$

## Discussion

Here we show that the cultural context in which a person is raised influences which objects they find memorable. It appears that Westerns tend to remember man-made objects better than things from nature, with our Chinese participants not showing such an asymmetry. This is consistent with prior research on cultural differences in memory. For instance, Westerns are more likely to remember salient foreground objects, while Easterners attend more to background context (Chua et al., 2005; Nisbett & Miyamoto, 2005). Bilingual studies have shown that people recall early-life events more vividly in their native language, implicating language and culture in memory formation (Marian & Fausey, 2006; Marian & Neisser, 2000; Marsh et al., 2015; Schrauf & Rubin, 2000). These lines of evidence suggest that cultural and linguistic contexts leave a lasting imprint on how information is encoded and retrieved.

Previous findings show high consistency in memorability across individuals, and even across species (Bainbridge et al., 2013; Isola et al., 2014; Khosla et al., 2015; Mancas & Le Meur, 2013; Jaegle et al., 2019). However, our data diverge from the hypothesis that memorability is entirely immune to individual experience. We demonstrate that cultural experience modulates how object features contribute to memorability: Western participants relied more heavily on physical properties (e.g., Metal/artificial and Tool/hammer), whereas Chinese participants incorporated both life-related features (e.g., Plant/green and Animal/organic) and physical properties. This suggests that while memorability has a strong intrinsic component, it can be systematically shaped by long-term environmental exposure. Our findings are a challenge to bottom-up accounts of memorability which predict that what is memorable is determined by aspects of the visual system that are shared across all humans (DiCarlo et al., 2012; Tai et al., 2017).

Our results further speak to the importance of the pre-training of long-term memory networks. According to computational models, new types of information should face less interference if they differ from previously stored content (Polyn et al., 2009; Shiffrin & Steyvers, 1997). However, we observed the opposite: participants showed expertise-driven memory advantages. American participants excelled at recalling tools and artifacts, while Chinese participants showed superior memory for natural objects. This likely reflects culturally distinct environmental exposures. Given that our participants were young adults, billions of objects would have already been viewed by the time they participated in our experiment (Huber, Geirhos et al. 2023), and these views shape the memory system in which the new information is received.

In sum, our cross-cultural study reveals how culturally scaffolded learning environments shape the memorability of novel stimuli. While memorability remains broadly consistent across humans—supporting its grounding in stimulus-driven properties—we identified culture-specific components that reflect differential weighting of features during storage. These findings reconcile universal constraints of visual processing with experience-dependent cognitive specialization.

## Acknowledgements

The work was funded by the National Science and Technology Innovation 2030 Major Program (2021ZD0200534) and the Natural Science Foundation of Guangdong grant (2023A1515012789) to BW; the National Science Foundation (BCS-2147064), and the National Institutes of Health (P30-EY08126) to GW. The code and behavioral data can be accessed through https://github.com/Haoryuchen/Prior-experience-on-memorability.

## References

Bainbridge, W. A. (2017). The memorability of people: Intrinsic memorability across transformations of a person’s face. *Journal of Experimental Psychology: Learning*, Memory, and Cognition, 43(5), 706–716. 10.1037/xlm0000339

Bainbridge, W. A. (2020). The resiliency of image memorability: A predictor of memory separate from attention and priming. Neuropsychologia, 141, 107408. 10.1016/j.neuropsychologia.2020.107408

Bainbridge, W. A., Isola, P., & Oliva, A. (2013). The intrinsic memorability of face photographs. Journal of Experimental Psychology. General, 142(4), 1323– 1334. 10.1037/a0033872

Chua, H. F., Boland, J. E., & Nisbett, R. E. (2005). Cultural variation in eye movements during scene perception. Proceedings of the National Academy of Sciences, 102(35), 12629–12633. 10.1073/pnas.0506162102

Connerton, P. (1989). How Societies Remember. Cambridge University Press.

DiCarlo, J. J., Zoccolan, D., & Rust, N. C. (2012). How Does the Brain Solve Visual Object Recognition? Neuron, 73(3), 415–434. 10.1016/j.neuron.2012.01.010

Hebart, M. N., Zheng, C. Y., Pereira, F., & Baker, C. I. (2020). Revealing the multidimensional mental representations of natural objects underlying human similarity judgements. Nature Human Behaviour, 4(11), 1173–1185. 10.1038/s41562-020-00951-3

Heersmink, R. (2023). Materialised Identities: Cultural Identity, Collective Memory, and Artifacts. Review of Philosophy and Psychology, 14(1), 249–265. 10.1007/s13164-021-00570-5

Hobsbawm, E., & Ranger, T. (Eds.). (2012). The Invention of Tradition. Cambridge University Press. 10.1017/CBO9781107295636

Huber, L. S., Geirhos, R., & Wichmann, F. A. (2023). The developmental trajectory of object recognition robustness: Children are like small adults but unlike big deep neural networks. Journal of Vision, 23(7), 4. 10.1167/jov.23.7.4

Isola, P., Xiao, J., Parikh, D., Torralba, A., & Oliva, A. (2014). What Makes a Photograph Memorable? IEEE Transactions on Pattern Analysis and Machine Intelligence, 36(7), 1469–1482. 10.1109/TPAMI.2013.200

Jaegle, A., Mehrpour, V., Mohsenzadeh, Y., Meyer, T., Oliva, A., & Rust, N. (2019). Population response magnitude variation in inferotemporal cortex predicts image memorability. eLife, 8, e47596. 10.7554/eLife.47596

Khosla, A., Raju, A. S., Torralba, A., & Oliva, A. (2015). Understanding and Predicting Image Memorability at a Large Scale. 2015 IEEE International Conference on Computer Vision (ICCV), 2390–2398. 10.1109/ICCV.2015.275

Kramer, M. A., Hebart, M. N., Baker, C. I., & Bainbridge, W. A. (2023). The features underlying the memorability of objects. Science Advances, 9(17), eadd2981. 10.1126/sciadv.add2981

Madan, C. R. (2021). Exploring word memorability: How well do different word properties explain item free-recall probability? Psychonomic Bulletin & Review, 28(2), 583–595. 10.3758/s13423-020-01820-w

Mancas, M., & Le Meur, O. (2013). Memorability of natural scenes: The role of attention. 2013 IEEE International Conference on Image Processing, 196–200. 10.1109/ICIP.2013.6738041

Marian, V., & Fausey, C. M. (2006). Language-dependent memory in bilingual learning. Applied Cognitive Psychology, 20(8), 1025–1047. 10.1002/acp.1242

Marian, V., & Neisser, U. (2000). Language-dependent recall of autobiographical memories. Journal of Experimental Psychology. General, 129(3), 361–368. 10.1037//0096-3445.129.3.361

Marsh, B. U., Kanaya, T., & Pezdek, K. (2015). The language dependent recall effect influences the number of items recalled in autobiographical memory reports. Journal of Cognitive Psychology, 27(7), 829–843. 10.1080/20445911.2015.1046876

Nisbett, R. E., & Miyamoto, Y. (2005). The influence of culture: Holistic versus analytic perception. Trends in Cognitive Sciences, 9(10), 467–473. 10.1016/j.tics.2005.08.004

Polyn, S. M., Norman, K. A., & Kahana, M. J. (2009). Task context and organization in free recall. Neuropsychologia, 47(11), 2158–2163. 10.1016/j.neuropsychologia.2009.02.013

Revsine, C., Goldberg, E., & Bainbridge, W. A. (2025). The memorability of voices is predictable and consistent across listeners. Nature Human Behaviour, 9(4), Article 4. 10.1038/s41562-025-02112-w

Saito, J. M., Kolisnyk, M., & Fukuda, K. (2023). Judgments of learning reveal conscious access to stimulus memorability. Psychonomic Bulletin & Review, 30(1), 317–330. 10.3758/s13423-022-02166-1

Schrauf, R. W., & Rubin, D. C. (2000). Internal languages of retrieval: The bilingual encoding of memories for the personal past. Memory & Cognition, 28(4), 616– 623. 10.3758/BF03201251

Shiffrin, R. M., & Steyvers, M. (1997). A model for recognition memory: REM— retrieving effectively from memory. Psychonomic Bulletin & Review, 4(2), 145–166. 10.3758/BF03209391

Tai, Y., Yang, J., Liu, X., & Xu, C. (2017). MemNet: A Persistent Memory Network for Image Restoration. 4539–4547. https://openaccess.thecvf.com/content_iccv_2017/html/Tai_MemNet_A_Persistent_ICCV_2017_paper.html

Xie, W., Bainbridge, W. A., Inati, S. K., Baker, C. I., & Zaghloul, K. A. (2020). Memorability of words in arbitrary verbal associations modulates memory retrieval in the anterior temporal lobe. Nature Human Behaviour, 4(9), 937– 948. 10.1038/s41562-020-0901-2

Zhao, C., Kim, J., Tang, T. H., Saito, J. M., & Fukuda, K. (2024). Deep neural network decodes aspects of stimulus-intrinsic memorability inaccessible to humans. Journal of Experimental Psychology. General, 153(4), 1131–1138. 10.1037/xge0001543

